# Metabolic adaptations leading to lignification in wheat roots under salinity stress

**DOI:** 10.1101/2023.06.08.544172

**Authors:** Bhagya M. Dissanayake, Christiana Staudinger, Kosala Ranathunge, Rana Munns, Thusitha W. Rupasinghe, Nicolas L. Taylor, A. Harvey Millar

**Affiliations:** School of Molecular Sciences, The University of Western Australia, 35 Stirling Highway, Crawley, Perth 6009, Australia; School of Biological Sciences, The University of Western Australia, 35 Stirling Highway, Crawley, Perth 6009, Australia; Institute of Agriculture, The University of Western Australia, 35 Stirling Highway, Crawley, Perth 6009, Australia; Institute of Agronomy, University of Natural Resources and Life Sciences, Konrad-Lorenz-Strasse 24, 3430 Tulln, Austria; Institute of Soil Research, University of Natural Resources and Life Sciences, Konrad-Lorenz-Strasse 24, 3430 Tulln, Austria; SCIEX, Mulgrave, Victoria, 3170, Australia

**Keywords:** bread wheat, tissue tolerance, salt exclusion, lignification, roots, proteomics

## Abstract

Analysis of salinity tolerance processes in wheat has focused on salt exclusion from shoots while root phenotypes have received limited attention. Here we consider the varying phenotypic response of four bread wheat varieties that differ in their type and degree of salt tolerance and consider in detail their molecular responses to salinity and changes in root cell wall lignification. These varieties were Westonia introgressed with *Nax1* and *Nax2* root sodium transporters (*HKT1;5-A*) that reduce Na^+^ accumulation in leaves, as well as the ‘tissue tolerant’ Portugese landrace Mocho de Espiga Branca that has a mutation in the homologous gene *HKT1;5-D* and has high Na^+^ concentration in leaves. These three varieties were compared with the more salt-sensitive cultivar Gladius. Through the use of root structural analysis, ion concentrations, as well as differential proteomics and targeted metabolomics we provide an integrated view of the wheat root response to salinity. We show different metabolic re-arrangements in energy conversion, primary metabolic machinery and phenylpropanoid pathway leading to monolignol production in a genotype and genotype by treatment dependent manner that alters the extent and localisation of root lignification which correlated with an improved capacity of wheat roots to cope better under salinity stress.

## Introduction

Soil salinity affects yields of a variety of crops across most continents of the world. Of the 230 million hectares of global irrigated land, 45 million hectares (19.5%) are affected by salt, while of the 1500 million hectares used for dryland agriculture, 32 million hectares (2.1%) are salt-affected soil (FAO, 2020). Bread wheat (*Triticum aestivum*) provides 20% of the total dietary requirements of humanity but the impact of salinity on its production can result in a yield loss of up to 60% in different parts of the world, contributing to food insecurity now and into the future (Dadshani et al. 2019). Most plants respond to salinity by exhibiting noticeable changes in root growth dynamics (Zou et al. 2022) and root system architecture (Shelden and Munns 2023). The wheat root system responds to saline conditions by an overall decrease in the length of the seminal (axile) roots and increase in the number of branch (lateral) roots (Rahnama et al. 2011). While much of the emphasis in salinity tolerance research has been placed on exclusion of salt Na^+^ and Cl^-^ from shoots (Moller et al. 2009; Wu and Li 2019) recent studies in different crops have also drawn a link between maintenance of plant salt tolerance and cell wall remodelling in roots. Changes in secondary cell wall composition including increase in lignification in roots under salinity stress has been reported in maize (Oliveira et al. 2020), soybean (Neves et al. 2010), barley (Ho et al. 2020), wheat (Jbir et al. 2001), and Arabidopsis (Reyt et al. 2021).

Radial and transverse primary cell walls deposit Casparian bands in the endo- and exodermis of roots (Steudle and Peterson 1998) and secondary cell walls, provides rigidity, and reducing transport of solutes and ions through the cell walls (apoplast) but promoting mineral transport through the cell-to-cell pathway of plants (Ranathunge and Schreiber 2011). Biosynthesis of lignin in plants occurs through substrates derived from primary metabolism entering the phenylpropanoid pathway (Weng and Chapple 2010). This pathway starts with the deamination of the proteinogenic amino acid phenylalanine by phenylalanine ammonia-lyase (PAL) to produce cinnamate, which is then converted to generate the three main monoliognols, p-coumaryl alcohol, coniferyl alcohol and sinapyl alcohol (Weng and Chapple 2010). During the last step, cell wall bound peroxidases and laccases catalyse the oxidative polymerisation of monolignols to produce lignin polymers (Neves et al. 2010). During recent years, the understanding of anatomical, gene expression and metabolic level changes in relation to root lignification has increased. Through the use of integrative multi-omics approaches Ho et al., 2020 reported that salt-induced modulation in apoplastic flow was linked to lignification and leading to salt tolerance in barley. Increased gene expression of genes including *CCoAOMT1*, *4CL1*, *4CL2*, *COMT*, *PAL1*, *PAL2*, and *AtPrx52* involved in the lignin biosynthesis pathway was observed in Arabidopsis roots under salinity stress in Arabidopsis roots, and a showed mutation in the *CCoAOMT1* gene leads to hypersensitivity to salinity (Chun et al. 2019). Further, the overexpression of superoxide dismutase (*SOD*) and ascorbate peroxidase (*APX*) enhanced the salt tolerance of transgenic *Arabidopsis thaliana* by increased oxidative polymerisation of monomers to accumulate lignin (Shafi et al. 2015). The study of Duan et al. (2020) provided further evidence of increased lignification in roots associated with salt tolerance in Arabidopsis, as did the study of wheat by Jbir et al. (2001) which showed histological and biochemical changes associated with differential lignin accumulation in bread wheat var. Tanit compared to durum wheat var. Ben Béchir and proposed the potential role of root lignification in maintaining salt tolerance in bread wheat. However, these studies have not directly addressed whether changes in upstream components of metabolism and biosynthesis in roots under saline conditions shift towards active lignification of secondary cell walls and the consequences of enhanced lignification on root growth and development in relation to salt tolerance.

In this study we have assessed root lignification responses of four different bread wheat varieties that differ in their mechanism and degree of saline tolerance. In bread wheat, salt tolerance is usually associated with low Na^+^ transport to the shoot and K^+^/Na^+^ discrimination (Na^+^ exclusion) (Munns et al. 2006). However recently, a Portugese landrace Mocho de Espiga Branca was identified by Borjigin et al. (2020) being “tissue tolerant” as it accumulates significantly higher Na^+^ concentrations while maintaining similar level of salt tolerance to Australian commercial bread wheat cultivars, such as Gladius. These four varieties, Gladius (a commercial bread wheat cultivar), Mocho de Espiga Branca (a tissue tolerant Portugese landrace), Westonia *Nax1* and Westonia *Nax2* (breeding lines introgressed with *Nax1* and *Nax2* root sodium transporters (*HKT1;5-A*) that reduce Na^+^ accumulation in leaves; James et al. 2011) were compared in terms of their anatomical, physiological, proteomic and lignin biosynthesis responses. The ‘tissue tolerant’ Portugese landrace Mocho de Espiga Branca has a mutation in the homologous gene *HKT1;5-D* and has high Na^+^ concentration in leaves yet retains growth in saline conditions (Borjigin et al., 2020). Our study provides new insights into metabolic re-arrangements in energy conversion and primary metabolic machinery in a genotype and genotype by treatment dependent manner that enhances the extent of lignification to improves the capacity of wheat roots to cope better under salinity stress.

## Materials and Methods

### Plant materials

Gladius and Mocho de Espiga Branca (Mocho) were kindly provided by A. Prof. Stuart Roy of the University of Adelaide and Westonia *Nax1* and Westonia *Nax2* were provided by CSIRO Agriculture and Food, Canberra. The source of the *Nax* genes was Australian bread wheat cultivar Westonia-derived BC_3_F_3_ fixed lines 5907 (*Nax1*) and 5924 (*Nax2*) respectively, the development of which was described previously by James et al. (2011).

### Plant growth in hydroponics

Mocho, Gladius, Westonia *Nax1* and Westonia *Nax2* were grown in half strength Hoagland solution in aerated hydroponics in 8 tanks of size 40x30x25 cm. Until the emergence of the third leaf, when application of salt began, tanks were in a growth chamber with 16 h/8 h light/dark cycle, and light intensity was maintained at 500 µmol m^-2^ s^-1^ at 28/22°C day/night temperature and 65% relative humidity. NaCl was added to the nutrient solution in 25 mM increments at 09:00 h and 16:00 h every day for 3 days to reach a final NaCl concentration of 150 mM. Each tank contained all four varieties, four tanks were maintained for each treatment (4 biological replicates). The initial 2 mM CaCl_2_ concentration of the half strength Hoagland solution was increased to 10 mM CaCl_2_ in the salt treatments. Four biological replicate plants of each variety for each treatment were harvested after 3 days and 6 days after reaching the final salt concentration.

### Measurement of plant growth parameters and ion concentrations

The rate of photosynthesis of the third leaf was measured by portable gas exchange system LI-6400 XT (LI-COR Biosciences, Lincoln, NE, USA) with microclimatic conditions adjusted in the leaf chamber to be similar with the plant growth environment. The light irradiation in the leaf chamber was set at 500 μmol m^−2^ s^−1^, leaf chamber temperature was stable at 24 °C and the CO_2_ reference concentration was 400 μmol mol^−1^. After this, the chlorophyll content was measured using a SPAD meter (Konica Minolta, Tokyo). To obtain the root length, roots were placed in a glass tray and scanned at 600 dpi using a desktop scanner (Epson Expression Scan 1680, Epson America Inc., Long Beach, CA, USA). The obtained images were analysed for root length and average root diameter using WinRHIZO 2008 (model Pro, version 2, Regent Instruments, Canada). After these measurements, roots and shoots were separated and dried in an oven at 70 °C for 5 days. (All the measurements were done in four biological replicates).

Dried root tissues and the fully expanded 3^rd^ leaf tissues were ground and digested with nitric acid (70% v/v) then with concentrated (70%) perchloric acid (Zarcinas et al. 1987). The Na^+^ and K^+^ concentrations were then quantified using inductively coupled plasma mass spectrometry (ICP-OES, Model 5300DV, Perkin Elmer, Shelton, CT, USA).

### Determination of root zones and lignin staining of roots

Healthy seminal roots without injuries were collected from three plants of each variety (Gladius, Mocho, Westonia *Nax1* and Westonia *Nax2*) grown under control and salt stress (150 mM NaCl) for 6 days. Roots were divided into two zones at the point where laterals were formed. Zone-I was the younger segment which included the actively growing root tip; this zone did not have lateral roots. Zone – II was the mature half or basal part of the root which included the lateral roots. The length of Zone-I and Zone-II varied according to the treatment. The average lengths of Zone-I and Zone-II were 13-15 cm, and 17-22 cm, respectively for all varieties under control conditions. Whereas the average lengths of Zone-I and Zone–II were 4-6 cm and 20-25 cm, respectively for all varieties under salt treatment. Zone-I was used for the anatomical studies where the structural modifications of the root took place. Since plants had different root lengths depending on the treatment, freehand cross-sections were made at 25, 50 and 75 % from the total length of Zone-I. These distances were measured from the root tip. This guaranteed that roots with the same percentage distance from the tip had the same age from both control and salt treatment, irrespective of their varied lengths. All root length measurements were started from the tip. To detect lignin in the cell walls of Zone-I, freehand cross-sections of roots were stained for several minutes with the Mäule reaction at room temperature (Ranathunge and Schreiber 2011). Lignified tissues were stained orange-red, whereas the rest tissues remained unstained under white light. Un-stained cross sections were viewed under Ultra-violet (UV) autofluorescence.

### Extraction of non-polymerised lignin monomers and intermediates

Briefly, 100 mg of roots was extracted in 75% methanol (v/v) (at 10 µL mg FW−1) for 2 h at 65 °C, and supernatant was collected after centrifugation for 20 min at 16,000 g. To concentrate the samples, 500 µL of supernatant was dried in speed-vac and then re-dissolved in 50 µL 50% methanol (v/v) and transferred to a standard HPLC vial. Subsequently, 10 μL was injected into the HPLC/MS/MS system for analysis.

### Targeted LC-MS to quantify phenylpropanoid pathway intermediates and monolignols

Targeted LC-MS analysis was carried out on a Sciex QTrap 6500+ triple quadrupole mass spectrometer from AB Sciex (Redwood City, CA, USA) coupled to Shimadzu Nexera LC-30AD UPLC system from Shimadzu (Columbia, MD) comprised of a vacuum degasser, quaternary pump, thermostat controlled auto-sampler and column oven. Chromatographic separations were performed on an Agilent InfinityLab Poroshell 120 HILIC-Z column (2.7 μm, 2.1 mm × 150 mm, Agilent Technologies, Santa Clara, CA) at a column temperature of 50°C and a flow rate of 0.8 mL/min. A linear gradient of aqueous solvent A (0.1% formic acid in H_2_O) and organic solvent B (0.1% formic acid in acetonitrile) were used as follows: 90% solvent B for 2.8 min, 90-80% solvent B over 0.2 min, 80-75% solvent B over 1.5 min, 75 – 50% solvent B over 0.5 min, 50-30% solvent B over 0.5 min, and hold at 30% solvent B over 1.5 min, return to 90% solvent B over 0.5 min, and equilibrate for 2.5 min at 90% solvent B resulting in a total run time for 10 min.

Lignin profiling was performed using QTrap 6500+ triple quadrupole with source parameters for the MS were set as follows: For the positive mode, curtain gas flow rate, 25 l/h; ion source voltage, 4.5 kV; desolvation temperature, 600 K; ion source gas 1 (GS1), 40 l/h; ion source gas 2 (GS2), 60 l/h. For the negative mode, curtain gas flow rate, 25 l/h; ion source voltage, -4.5 kV; desolvation temperature, 600 K; ion source gas 1 (GS1), 40 l/h; ion source gas 2 (GS2), 60 l/h. ESI parameters for both negative and positive mode were set up for delustering potential (DP), entrance potential (EP), collision energy (CE) and cell exit potential (CXP) for each metabolite were shown in the Supplementary Data 1 . Data has been processed using SCIEX OS software (version 2.2) before exporting the peak area to an excel sheet for further calculations.

### Cell wall isolation of wheat roots

To quantify polymerised lignin in cell walls which makes the transport barrier through the apoplast/cell walls, roots of Gladius, Mocho, Westonia *Nax1* and Westonia *Nax2* grown under control and salt treatment conditions (150 mM NaCl) for 6 days were used. Zone-I and Zone-II root segments were incubated in an enzymatic solution containing 1 % (v/v) cellulase and 1 % (v/v) pectinase (Chem Supply Pty Ltd, Gillman, SA, Australia) for 4 weeks. Isolated wall samples were thoroughly extracted in chloroform and methanol (1:1; v/v) for 2 days and were dried and stored over silica gel. Five milligrams of dried wall samples were used to analyse polymerised lignin (Johnson et al. 1961). Sample digestion with acetyl bromide and addition of 2 M NaOH and glacial acetic acid to hydrolyse excess acetyl bromide were carried out as described previously by (Ranathunge and Schreiber 2011). To destroy organic bromides, 7.5 M hydroxylamine-HCl was added to the samples. The absorbance at 280 nm of each sample was measured with a spectrophotometer (MULTISKAN GO, Thermo Fisher Scientific, Vantaa, Finland). Four biological replicates were used for each genotype and treatment.

### Root protein extraction and digestion

A 150 mg sample of root tissue was snap frozen and ground with glass beads to a fine powder, then 300 μL of extraction solution (125 mM Tris-HCl pH 7.5, 7% (w/v) sodium dodecyl sulfate (SDS), 0.5% (w/v) PVP40 with Roche protease inhibitor cocktail (Roche) added at 1 tablet per 50 mL) was added. The protein fraction was collected by a chloroform/methanol extraction (Wessel and Flugge 1984), the pellet obtained was washed twice with 80% (v/v) acetone. Samples were resuspended in 60 µL freshly prepared resuspension buffer (7 M urea, 2 M thiourea, 50 mM NH_4_HCO_3_ and 10 mM DTT). The protein concentration was determined by the addition of Bradford reagent (Thermo scientific) and spectrophotometric measurement at a wavelength of 595 nm using bovine serum albumin as a standard.

One hundred micrograms of protein from each sample was treated with 25 mM iodoacetamide for 30 minutes in the dark at room temperature. The sample was then diluted (1:10) with 20 mM NH_4_HCO_3_. Trypsin was prepared by adding 25 μL of 0.01% (w/v) trifluoroacetic acid (TFA). For protein digestion, trypsin was added to each sample at a trypsin:protein ratio of 1:25. The samples were then incubated at 37° C overnight and the next morning the samples were acidified to 1% (v/v) with formic acid in order to stop the enzymatic reaction.

Peptide samples were purified using Silica C18 Macrospin columns (The Nest Group). After each step of the purification, solid phase extraction columns were centrifuged for 2 min at 1000 x g at room temperature. Before loading the cartridge with peptides it was charged by addition of 200 μL of 100% (v/v) methanol, then the column was equilibrated with 200 µL 5%(v/v) acetonitrile (ACN), 0.1% (v/v) formic acid (FA). After loading the samples, the column was washed with 5% (v/v) ACN, 0.1% (v/v) FA twice followed by the elution of peptides twice with 70% (v/v) ACN, 0.1% (v/v) FA. The elute was dried under vacuum and resuspended in 5% (v/v) ACN, 0.1% (v/v) FA to a final concentration of 1 μg/μL.

### Quantitative untargeted peptide mass spectrometry

Root protein samples were injected (0.5 µg protein) into an online nanoflow (300 nL / min) capillary column (Picofrit with 10μm tip opening / 75μm diameter, New Objective, PF360–75-10-N-5) packed in-house with 15 cm C18 silica material (3 µm; Dr. Maisch GmbH) connected to a ThermoFisher Exploris 480, in-line with Dionex Ultimate 3000 series UHPLC. Spray voltage were set to 1.9 kV, and heated capillary temperature at 250 °C. For DDA experiments full MS resolutions were set to 120,000 at m/z 200 and full MS AGC target was 300% with an IT of 40 ms. Mass range was set to 350–1700. AGC target value for fragment spectra was set at 100% with a resolution of 15,000 and injection times of 28 ms. Intensity threshold was kept at 2E4. Isolation width was set at 1.4 m/z. Normalized collision energy was set at 30%. The acquired spectra were then searched against Uniprot reference proteome (UP000019116) of Triticum aestivum cv. Chinese Spring using MaxQuant version 1.6.17.0. The searching parameters were as follows: 0.5 Da mass tolerance for peptides and 0.01 Da mass tolerance for fragments, carbamidomethyl-cysteine as fixed modification, oxidation of methionine and protein-N-acetylation as variable modifications. The shot-gun proteomics data have been deposited to the ProteomeXchange Consortium (http://proteomecentral.proteomeexchange.org) via PRIDE partner repository Vizcaino et al. (2013) with the dataset identifier PXD041669.

### Differential protein abundance (DAP) analysis

MaxQuant output files were analysed though R software (Version 3.6.1) using the Bioconductor (DEP 1.1.4 package) (package for differentially expressed which is specifically designed to analyse mass spectrometry based proteomic data and uses MaxQuant output directly as the input files. Among the 6120 total proteins detected by Orbitrap, 3784 proteins were retained after filtering the missing values through a stringent filter where proteins which are identified in at least 70% of the total number of samples within one variety tested for both time points were retained. Background correction and normalization was performed by variance stabilising transformation (vsn) method. Missing values were imputed through MinProb approach followed by the differential enrichment analysis conducted for comparison of salt treatment vs. control conditions for all wheat varieties. Proteins which have higher fold changes than log2 (1.5) and p values less than 0.05 were considered as differentially expressed. Hierarchical clustering was performed for the proteins which were shared by all four varieties at each time point.

### 2 WAY ANOVA analysis of quantitative proteomics data

2-way ANOVA was conducted on all the protein abundances to identify, genotypic response, treatment response and the genotype x treatment interaction under salt stress conditions. 2-way ANOVA was conducted for the imputed, normalized dataset by R (version 3.6.1) with the use of “rstatix” package which provides a pipeline of functions for statistical analyses. Proteins found as significant (p<0.05) for the genotype, treatment and genotype x treatment interaction by 2-way ANOVA were then subjected to Tuckey’s post hoc test to obtain adjusted p values for all possible genotype combinations including Mocho-Gladius, Westonia Nax1-Gladius, Westonia *Nax2*-Gladius, Westonia *Nax1*-Mocho, Westonia *Nax2*-Mocho, Westonia *Nax1*-Westonia Nax2. Tuckey’s post hoc test was conducted to obtain proteins significant for the most important genotype x treatment interaction combinations which includes Mocho:C-Gladius:C, *Nax1*:C-Gladius:C, *Nax2*:C-Gadius:C, Mocho:S-Gladius:S, *Nax1*:S-Gladius:S, *Nax2*:S-Gladius:S.

## Results

### The impact of salinity stress on wheat root growth and physiology

In order to determine the effect of salt stress on growth and physiology of Gladius, Mocho, Westonia *Nax1* and Westonia *Nax2* plants, the dry weights of shoots and roots were measured 3 and 6 days after reaching the final NaCl concentration of 150 mM in the hydroponic growth media. After 3 days there were few statistically significant differences in growth characteristics between the control and salt treatment in any variety (Supplementary Figure 1). However, after 6 days, all growth characteristics were affected by the salinity treatment (Figure 1). The root dry weight and shoot dry weight of Gladius, Mocho and Westonia Nax1 was decreased by salt, but root DW of Westonia *Nax2* was unaffected (Figure 1A). A significant decrease in total root length was observed in salt treated roots of Mocho and Gladius, Westonia *Nax1* but not in Westonia *Nax2* (Figure 1B1). Root length of Gladius was the most affected and showed a significant decrease in root length in all measured root diameter classes (Figure 1B2). Mocho and Westonia *Nax1* showed decreases in the smaller diameter roots but did not show significant changes in roots of the larger diameter classes, 1.0-1.5 mm and 1.52.0 mm (Figure 1B2). In Westonia *Nax2* the length of roots belonging to the larger diameter classes increased under salt exposure and was significantly greater than the other varieties (Figure 1B2) which indicates that Westonia *Nax2* produces thicker roots under salt stress.

In terms of ion concentrations in roots, all the varieties showed approximately a 2-fold increase in Na^+^ concentration after 3 d (Supplementary Figure 1C) and 6 d (Figure 1C) of salt stress. After 6 d of salt stress there were no significant differences between varieties (Figure 1C). The K^+^ concentration of roots in all varieties was reduced significantly after 3 d (Supplementary Figure 1D), but after 6 days there were no significant differences between treatments or between varieties (Figure 1D). Even though, salt treatment significantly reduced the K^+^/Na^+^ ratio, the K^+^/Na^+^ ratio was similar in all varieties at both times after salt addition (Supplementary Figure 1E, Figure 1E).

**Figure 1.**
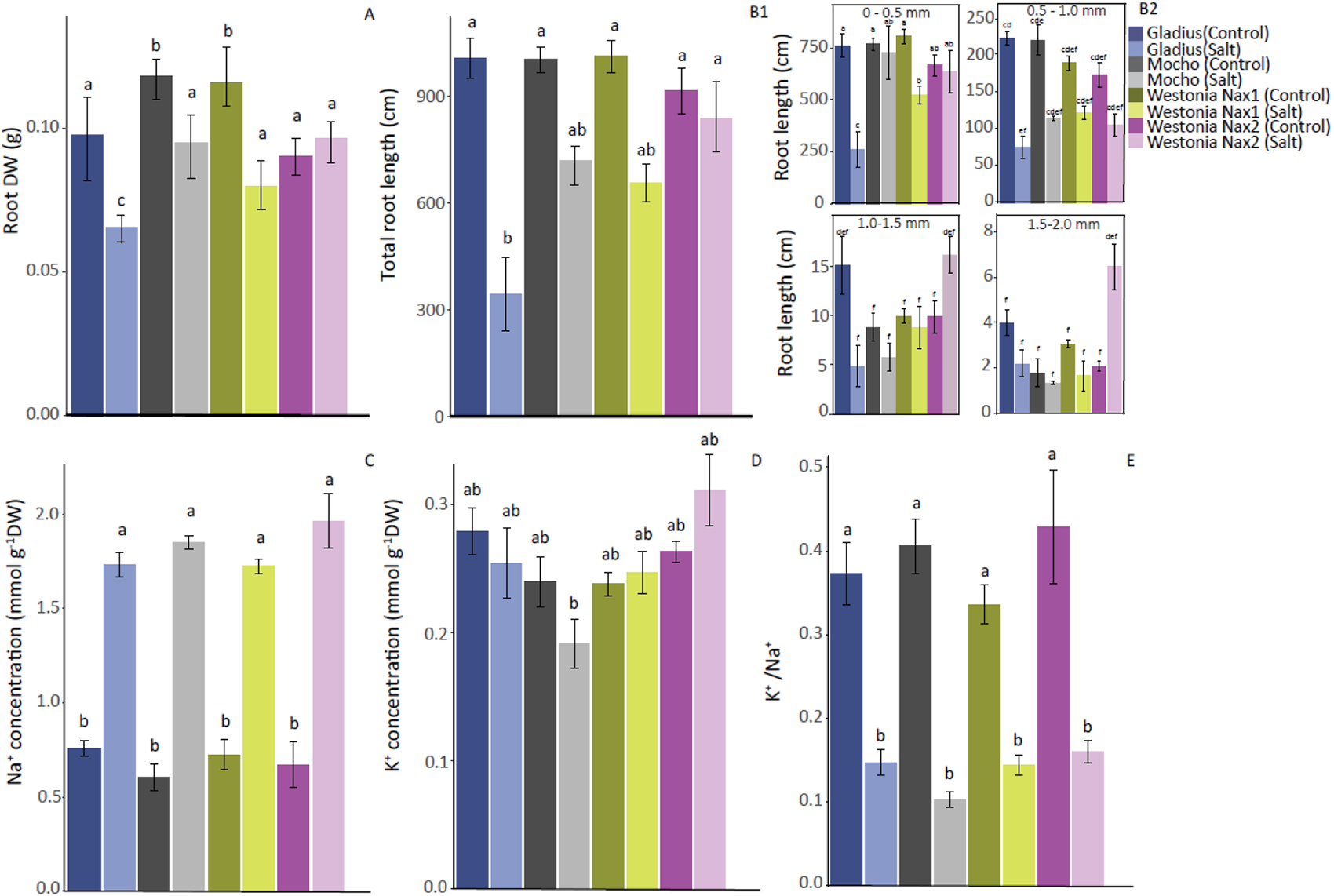
Differences in root growth and root ion concentrations in the four varieties after 6 days of exposure to 150 mM NaCl. (A) Root dry weight, (B1) Total root length, (B2) Root lengths that belong to different diameter classes, (C) Na^+^ ion concentration, (D) K^+^ ion concentration, (E) K^+^/Na^+^ ratio. Different letters denote significant differences obtained through Tukey’s HSD test, error bars indicate the standard error of mean, n=4.

Shoot dry weight of all four varieties reduced in response to salinity, however Mocho maintained the highest DW under salt stress (Figure 2A). Mocho accumulated 3-4-fold higher Na^+^ concentration compared to other varieties after salt stress was applied (Figure 2B). Westonia *Nax1* and *Nax2* maintained significantly lower Na^+^ concentrations compared to Gladius and Mocho (Figure 2B). The K^+^ concentration of all varieties was significantly lower compared with their respective controls (Figure 2C), but Westonia *Nax1* and *Nax2* maintained significantly higher K^+^/Na^+^ ratios (Figure 2D) compared to Gladius and Mocho which indicates the capability of roots containing *Nax1* and *Nax2* genes to discriminate K^+^ over Na^+^ under salt stress conditions and prevent it reaching the shoots.

The salt treatment significantly reduced the chlorophyll content of all varieties (Figure 2E). Rate of photosynthesis was also reduced in all varieties under salt treatment, but Westonia *Nax* lines maintained higher rate compared to other varieties under salt treatment (Figure 2F).

Overall, the greatest decrease in growth characteristics was shown in Gladius, with Mocho the next most affected. Westonia *Nax2* performed best of the four with the least decrease in root dry weight and in rate of photosynthesis.

**Figure 2.**
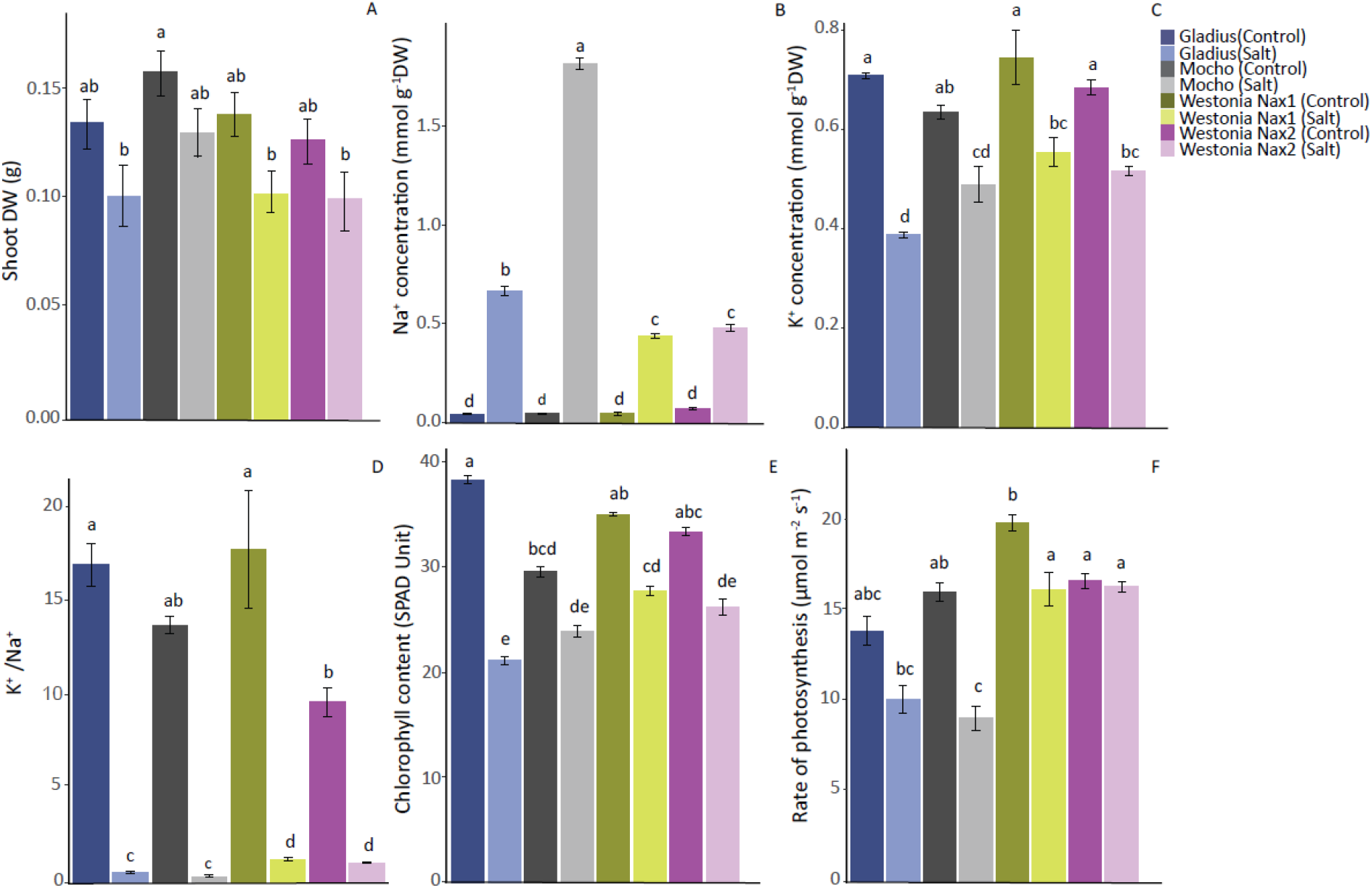
Differences in shoot growth, Na^+^ and K^+^ ion concentrations, photosynthetic rate after 6 days of exposure to 150 mM NaCl. (A) Shoot Dry weight, (B) Na^+^ ion concentration, (D) K^+^ ion concentration, (E) The K^+^/Na^+^ ratio. Different letters denote significant differences obtained through Tukey’s HSD test, error bars indicate the standard error of mean, n=4.

### Differences in wheat root lignification under salt treatment

Lignin in the cell walls of root cross sections was detected by an orange/red staining using the Mäule reaction. In control, more lignification was observed through bright orange/red staining in early metaxylem in the vascular cylinder (VC) of Mocho and Westonia *Nax* lines compared to Gladius that were the same age; 75% distance from the root tip of the Zone-I (Figure 3A-D, ∼11.25 cm from the root tip; Supplementary Figure 2A-D). More intense lignification in the cell walls of early and late metaxylem vessels was observed though dark red/pink staining in all four varieties under salt treatment. However, the intensity of the lignification of the stele was much higher in Westonia *Nax* lines compared to Gladius and Mocho. Except Gladius, most of the parenchymal cells surrounding the stele were lignified under salt treatment. Further, under salt treatment, the “U” shaped tertiary cell walls in the endodermis were present in all varieties (arrows in Figure 3A-D) but the number of cells without “U” shaped cell walls (passage cells) was greater in Mocho than other varieties (Figure 3B *vs* 3A,C,D). The endodermal “U” shaped tertiary cell walls were stained brighter orange (arrows) in roots of all varieties indicating the deposition of salinity-induced lignin into cell walls Figure 3A-D). Thickness of the “U” shaped endodermal cell walls were much higher in Westonia *Nax* lines followed by Gladius, while Mocho was the least. Moreover, under salt treated condition, the number of endodermal cells without “U” shaped walls (passage cells) was greater in Mocho than other varieties (Figure 3B). Even though histochemical staining showed less lignification in the VC of Gladius compared with other varieties (Supplementary Figure 2A-D), the total lignin amount of Gladius roots was not statistically different from the other varieties under control growth (Figure 3I). In agreement with the histochemical staining the salinity-induced lignin amounts in roots of Mocho, Westonia *Nax1* and Westonia *Nax2* were significantly higher than that of Gladius. Overall, roots of Gladius remained the least lignified among all four varieties under both control and salt treatment. Thickness of the endodermal “U” shaped cells were much higher in Westonia *Nax1* and Westonia *Nax2* followed by Gladius (Figure 3E-H), However, Mocho had the least thickness in endodermal cell walls with the highest number of passage cells compared to Gladius and Westonia *Nax* lines (Figure 3E-H).

**Figure 3.**
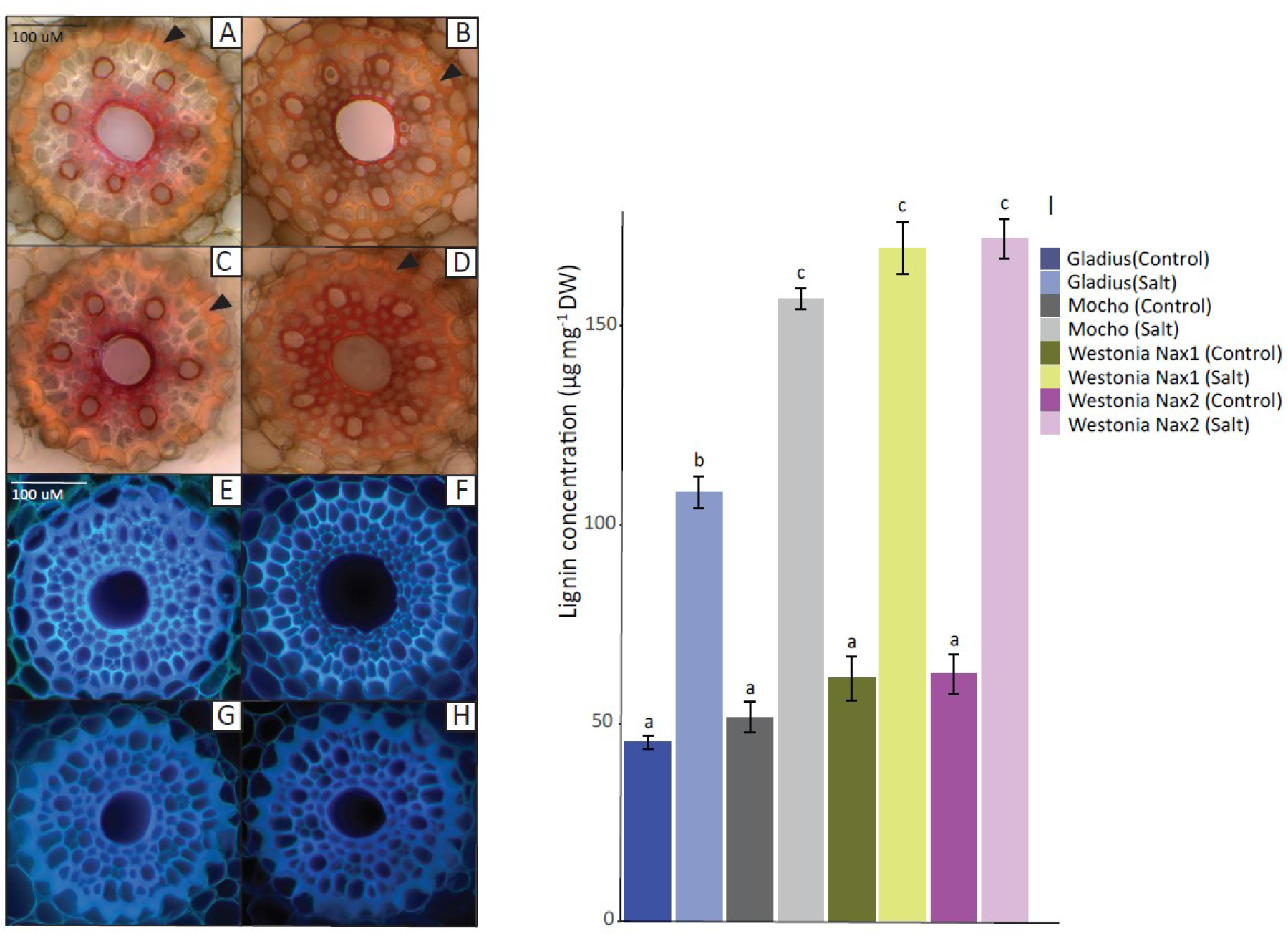
Comparison of lignin depositions and changes in the concentration of total polymerised lignin in roots. Lignin staining in the vascular cylinder (VC) and endodermis of roots gown under 150 mM NaCl for 6 days in an aerated hydroponic system (A–D); roots grown under 150 mM NaCl for 6 days in an aerated hydroponic system and observed under UV autofluorescence (E–H). At 75 % from the total length of Zone-I, orange/red staining in the early VC and endodermis of roots grown under salt treatment condition: (A) Faint orange staining in the endodermis and bright red staining in VC, (B) Orange staining in the endodermis and dark red staining in VC of Mocho, (C) Orange staining in the endodermis and dark red staining in VC of Westonia *Nax1*, and (D) Orange staining in the endodermis and dark red staining in VC of Westonia *Nax2*. (A-D) The “U” shaped tertiary cell walls in the endodermis with orange stains (arrows) in roots. At 75 % from the total length of Zone-I, un-stained cross sections of roots grown under roots grown under salt treatment condition: (E) Gladius, (F) Mocho, (G) Westonia *Nax1*, and (H) Westonia *Nax2*. Bars 100 µm. (I) The Total amounts of root lignin in roots from plants grown either in control or in 150 mM NaCl treatment concentrations after 6 days. Data are means ± SE (n = 3). Different letters denote significant differences obtained through Tukey’s HSD test at P ≤ 0.05 level.

### Quantitative changes in the wheat root proteome under salinity stress

In order to identify the salt responsive proteins in wheat roots among the 3784 proteins we could quantify in control and salt treated samples, a differentially abundant protein analysis (DAP) was conducted. After 6 days of salt stress 633, 688, 604 and 735 protein groups were differentially abundant in Gladius, Mocho, Westonia *Nax1* and Westonia *Nax2* respectively. One hundred and thirty-nine DAPs were common to all four varieties, whereas 188, 248, 210 and 313 DAPs were found to be unique to Gladius, Mocho, Westonia *Nax1* and Westonia *Nax2* respectively (Figure 4B). A number of protein responses were identified as being common across genotypes at both 3 days and 6 days indicating the generic wheat root response to salt stress. While the intensity of these generic responses for some proteins varies, the direction of the change in abundance was similar for all four varieties, consistent with the notion of it representing a generic wheat protein abundance response in roots to salt stress (Supplementary Figure 3 and Supplementary Figure 4). The majority of proteins which increased in abundance after 6 days of salt stress were found to be involved in stress and transport functional categories (Supplementary Figure 4). Most of the DAPs which showed a decrease in abundance at both time points were involved in protein synthesis and degradation and secondary metabolism (Supplementary Figure 3 and Supplementary Figure 4). This suggests that the machinery for protein metabolism in wheat roots in typically decreased by salt stress across genotypes.

When examining the number of DAPs, after 3 and 6 days of salt stress, a significant number of DAPs were observed in only one variety, indicating unique responses to salt stress (Figure 4B, Supplementary Figure 4). DAPs which were uniquely expressed in Westonia *Nax1* and Westonia *Nax2* included proteins involved in glycolysis, the TCA cycle and oxidative phosphorylation which indicates that the roots of these cultivars enhance the machinery of energy metabolism under salinity stress. In Westonia *Nax1* central metabolic enzymes (PEPC and enolase) and sugar degradation enzymes (fructokinase and hexokinase) (Supplementary Data 2) were increased in abundance, indicating an enhancement of glycolytic metabolism under salt stress (Figure 4A). Interestingly, in Westonia *Nax1* proteins involved in aromatic amino acid synthesis were found to significantly increased in abundance after salt exposure (e.g. 3-deoxy-D-arabino-heptulosonate 7-phosphate synthase, anthranilate phosphoribosyltransferase and tryptophan synthase) (Figure 4A, Supplementary Data 2). The observed increases in abundance of two 3-deoxy-D-arabino-heptulosonate 7-phosphate synthases indicates that the shikimate pathway is altered in the Westonia *Nax1* line during salt exposure. Several glycolytic enzymes such as enolase, glyceraldehyde 3-phosphate dehydrogenase (GAPDH), phosphofructokinase (PFK), glucose-6-phosphate isomerase and pyruvate kinase (PK) (Figure 4A, Supplementary Data 3) increased in abundance in Westonia *Nax2*. Increased abundance of these proteins likely indicates an increased investment in the glycolytic machinery. Increases in abundance of enzymes involved in oxidative phosphorylation (cytochrome c oxidase subunit 5B, ATP synthase gamma chain, ATP synthase delta-subunit and ATP synthase subunit d) (Figure 4A, Supplementary Data 3), the TCA cycle (acetyltransferase (E2) component of pyruvate dehydrogenase and isocitrate dehydrogenase) and those in glycolysis suggests an adaptation to increase ATP production under salt stress conditions. However, in Mocho such an increase in glycolytic machinery wasn’t observed, on the contrary the TCA cycle enzymes malate dehydrogenase, isocitrate dehydrogenase, pyruvate dehydrogenase (E2 subunit) and mitochondrial oxidative transport related external alternative NAD(P)H-ubiquinone oxidoreductase B2 were decreased in abundance under salt stress.

**Figure 4.**
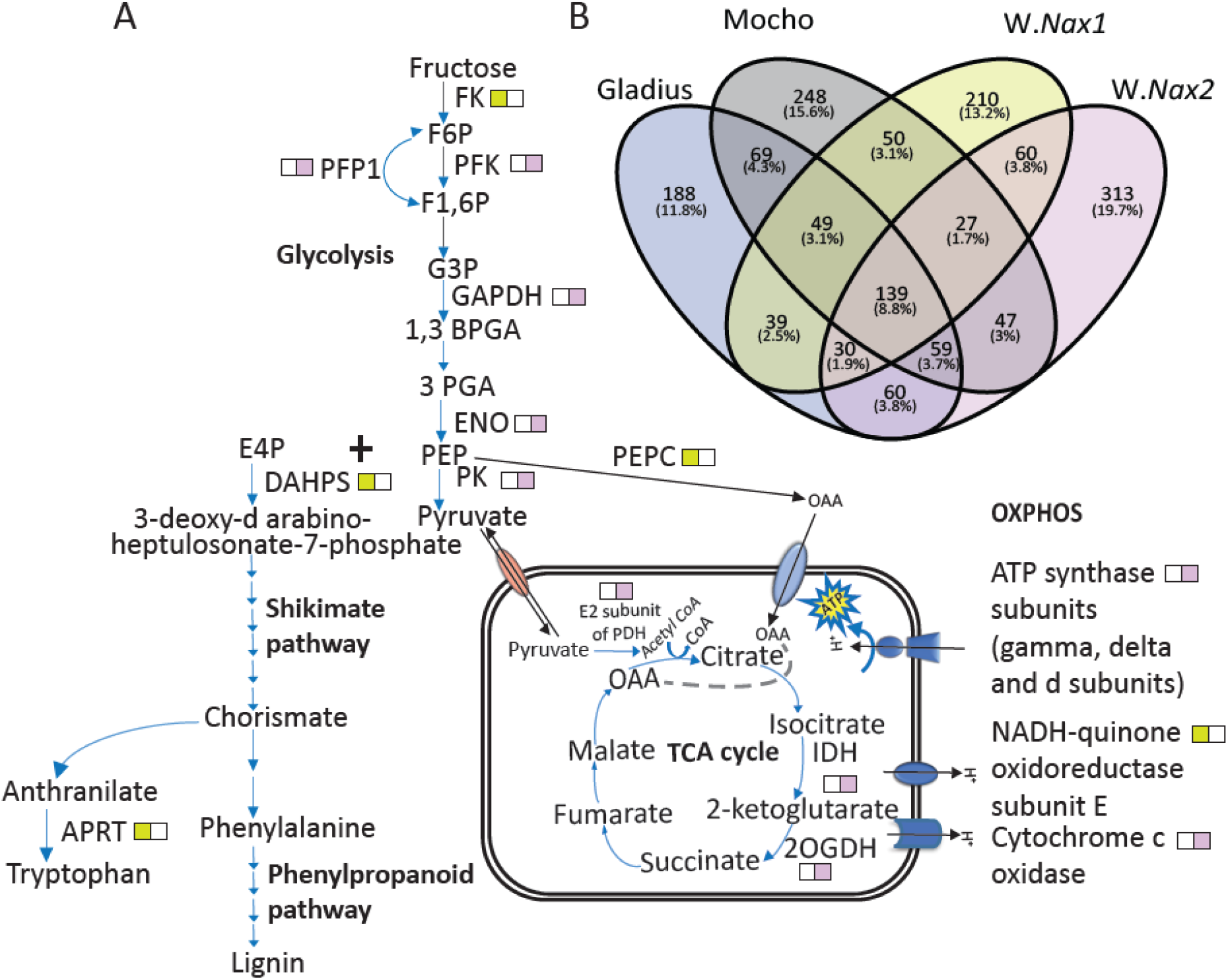
(A) DAPs which showed increased in abundance in W.*Nax1* (left box) DAPs which showed increased in abundance in W.*Nax2* (right box) after 6 days of exposure to 150mM NaCl treatment (B) Venn diagrams showing the overlap of DAP numbers and respective percentages of the total number of proteins identified. Colours in A are respective to the colours in B for W.*Nax1* and W.*Nax2*. Abbreviations: W.*Nax1*; Westonia *Nax1*, W.*Nax2*; Westonia *Nax2*.

### 2-Way ANOVA genotypic response

While the DAP analysis clearly highlighted root proteomic responses to salinity treatment, it did not readily provide information on constitutive differences between genotypes. To observe these genotypic differences, we investigated genotype x salinity treatment interactions of the root proteomes using 2-way ANOVA analysis. When comparing the three most salt tolerant to the least tolerant variety (i.e. Gladius), 160, 127, 186 proteins were found to be significantly different for the Mocho vs. Gladius, Westonia *Nax1* vs. Gladius, Westonia *Nax2* vs. Gladius comparisons, respectively. In the Mocho vs. Gladius comparison, 71 proteins were found be significantly more abundant in Mocho than in Gladius (Supplementary Data 4), whereas 89 proteins were found to be more abundant in Gladius compared to Mocho (Supplementary Data 4). 3-dehydroquinate synthase and phospho-2-dehydro-3-deoxyheptonate aldolase which are involved in the first and second steps of the shikimate pathway/chorismate pathway were constitutively higher in abundance in Mocho compared to Gladius (Supplementary Data 4). This could indicate that carbon flow is diverted to the shikimate, pathway during salt exposure to generate chorismate, a precursor to synthesise aromatic amino acids in Mocho. The lower abundance of enzymes such as isocitrate dehydrogenase, fumarate hydratase, 2-oxoglutarate dehydrogenase in roots of Mocho (Supplementary Data 4) may also indicate that the diversion of carbon flow through PEP to the shikimate pathway reduces the carbon flow to the TCA cycle in Mocho.

In the genotype comparison of both Westonia *Nax* lines with Gladius, 43 proteins were found to be significantly more abundant in Westonia *Nax* lines compared to Gladius, whereas 37 proteins were found to be more abundant in Gladius compared to Westonia *Nax* lines (Supplementary Data 5). A number of proteins involved in secondary metabolism including 3-ketoacyl-CoA thiolase, hydroxycinnamaldehyde dehydrogenase and cytochrome P450 family cinnamate 4-hydroxylase (C4H) were more abundant in Westonia *Nax* lines than Gladius. Moreover, cinnamoyl alcohol dehydrogenase (CAD) which catalyses a committed step in lignin biosynthesis were found to be more abundant in both Westonia *Nax* lines compared to Gladius (Supplementary Data 5).

Focusing on what was unique to Gladius, we observed a low abundance of pyruvate decarboxylase, and a higher abundance of malic enzyme and alanine aminotransferase compared to Mocho, W.*Nax1* and W.*Nax2* through genotype x treatment interaction of 2-Way ANOVA. This could indicate that Gladius undergoes a metabolic re-arrangement at the end of glycolysis to improve downstream pyruvate metabolism (Supplementary Data 6) rather than altering secondary metabolic pathways towards lignin biosynthesis.

### Targeted metabolic analysis of the phenylpropanoid pathway to monolignols

In order to verify if changes in protein abundance in lignin biosynthesis machinery in roots was altering lignin formation, a targeted MRM approach was used to quantify the phenylpropanoid pathway intermediates leading to the synthesis of monolignols. In general, many of the quantified phenylpropanoid pathway intermediates were more abundant in Mocho and Westonia *Nax* lines compared to Gladius under both control and salt treatment conditions (Figure 5). Most of the significant changes in abundance were observed under salt stress conditions compared to controls. All three monolignol subunits (coniferyl alcohol, sinapyl alcohol and p-coumaryl alcohol) were found to be significantly more abundant in Westonia *Nax2* under salt stress. While roots of Mocho also accumulated a significantly higher amount of coniferyl alcohol (Figure 5). Westonia *Nax1* also accumulated higher levels of all three monolignols, however the abundance changes were not significant. High abundance of phenylpropanoid pathway intermediates and monolignols in Mocho, Westonia *Nax1* and Westonia *Nax2* is consistent with a metabolic re-arrangement to obtain a higher capacity for root lignification under salinity stress.

**Figure 5.**
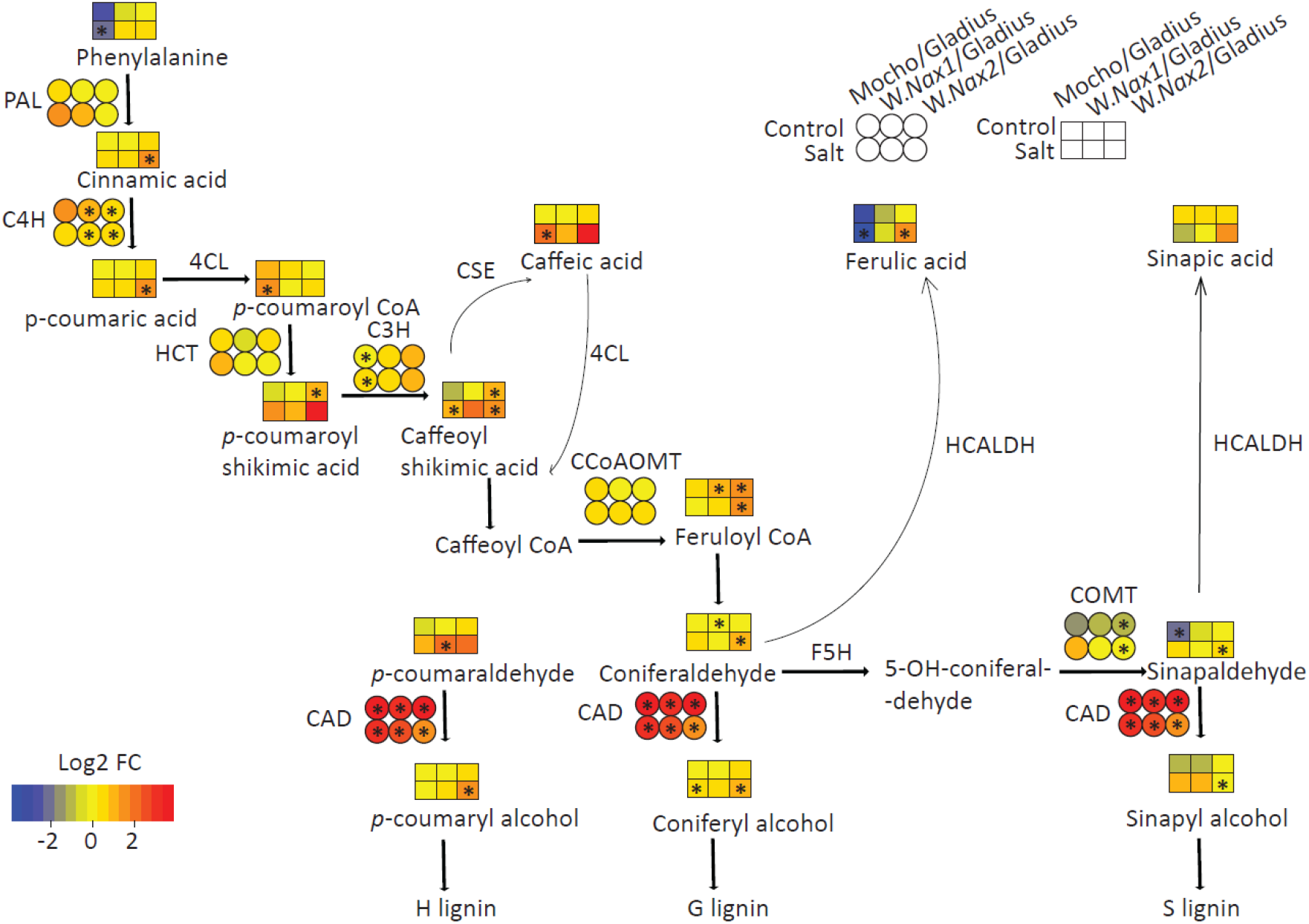
Diagram illustrating constitutive and induced protein and metabolite changes in roots of bread wheat varieties. Asterisks indicate significant differences (2-Way ANOVA followed by Tukey’s HSD test for proteins, t-test for pairwise analysis of metabolites; * P < 0.05, the protein abundances are illustrated in circles, metabolite abundances are illustrated in squares, n = 4) PAL-Phenylalanine ammonia lyase, HCALDH-Hydroxycinnamaldehyde dehydrogenase, CSE-Caffeoyl shikimate esterase, 4CL-4-coumarate:CoA ligase, C4H-Cinnamic acid 4 hydroxylase, C3H-p-coumaroyl shikimate 3ʹ hydroxylase, HCT-Hydroxycinnamoyl-coenzyme A shikimate:quinate hydroxycinnamoyl-transferase, CCoAOMT – Caffeoyl-CoA O methyltransferase, F5H-flavanone 5 hydroxylase, COMT – Caffeic acid/5-hydroxyferulic acid O-methyltransferase, CAD – Cinnamoyl alcohol dehydrogenase.

## Discussion

A range of studies have suggested the importance of root lignification to maintain salt tolerance in plants including Arabidopsis (Chun et al. 2019; Naseer et al. 2012), wheat (Jbir et al. 2001), barley (Ho et al. 2020), maize (Oliveira et al. 2020), soybean (Neves et al. 2010), Sophora alopecuroides (Zhu et al. 2021), and tomato (Sánchez-Aguayo et al. 2004). Lignin is involved during the first phase of the salt stress response which causes damage to the plant if the stress level is high, and the plant does not possess appropriate salt tolerant mechanisms (Cabane et al. 2012). However, the exact role which lignification plays to maintain salt tolerance of crops and how roots are enabled to increase their level of lignification is yet be understood.

So far we know that root lignification plays a role in maintaining firmness and of secondary cell walls (Ho et al. 2020; Zou et al. 2022), limiting the entry of Na^+^ by creating apoplastic barriers (Sexauer et al. 2021). The contribution of cell wall integrity to salt tolerance in plants has been well documented (Colin et al. 2023). Naseer et al. (2012) reported that deposition of lignin contributes to the mechanical strength of the cell wall of roots as well as protection of membrane integrity under salt stress. Chun et al. (2019) reported Arabidopsis root cells showed thicker cell walls and increased lignin concentration under high salt conditions indicating the importance of physical reinforcement of cell walls for long-term adaptation to salt stress.

Based on this, we propose that the more intense lignification observed in salt tolerant lines Mocho, Westonia *Nax1* and Westonia *Nax2* than in Gladius allows these varieties to cope better under salt stress through physical reinforcement of root cell walls. Further, Westonia *Nax* lines may control Na^+^ entry through forming effective root apoplastic barriers. Root growth reduction was observed in maize (Neumann et al. 1994) and soybean (Neves et al. 2010) by lignification of cell walls in the growing root tips. However, in our study, lignification of root tip cells did not occur. Therefore, we propose the lignification in salt tolerant lines is associated with sustained root growth by reinforcing against the unfavourable circumstances caused by salinity stress. This is in agreement with our physiology data in which Mocho, Westonia *Nax1* and Westonia *Nax2* maintained significantly higher root length compared to Gladius under salt stress. Similar observation was also reported by (Ho et al. 2020), in which the authors suggested that salt induced lignification sustained the root growth of salt tolerant barley cv. Clipper.

The root exodermis and endodermis form transport barriers by modifications of chemical compositions of cell wall (Schreiber et al. 1999). Generally, the primary endodermis is considered as the main apoplastic barrier for the passive movement of water and dissolved ions from the soil solution (Schreiber et al. 1999). Apoplastic barrier in endodermis forces the passively travelled ions to pass through the plasma membrane of living endodermal cells where specific transporters control the entry to the stele (Chen et al. 2011). In monocots like wheat, the final stage of the endodermis is characterised by extensive thickening of cell walls by deposition of cellulose oriented towards the stele rather than the cortex (Byrt et al. 2018). Consistent with this, extensively thickened and lignified cell walls in the endodermis were observed in both Westonia *Nax1* and Westonia *Nax2* under salt stress, which indicates development of a functioning endodermis under salt stress. In addition to the *Nax* transporters, the functional endodermal barrier could provide an additional capacity to restrict the entry of Na^+^ to the xylem which contributes to the reduced amount of Na^+^ levels observed in the shoots. With less number of “U” shaped cells within the endodermis and relatively thin walled cells in the endodermis could contribute to the additional Na^+^ entry to the root xylem which then gets accumulated within the shoots. Apart from the single nucleotide substitution in *TaHKT1;5-D* Na^+^ transporter in Mocho (Borjigin et al. 2020), the formation of less effective endodermal apoplastic barrier in roots will further enhances the entry of Na^+^ to the root xylem. In addition to that we observed lignification of pith parenchymal tissues under salinity. This interpretation of changes in lignification as a salt tolerance strategy is consistent with the report by (Jbir et al. 2001) that lignification was more pronounced in the VC of a salt tolerant variety than the salt sensitive variety. Increases in root lignification and in the number of lignified xylem vessels have also been reported in soybean (Neves et al. 2010) and tomato (Sánchez-Aguayo et al. 2004) in response to salinity. Similar lignification patterns have been observed in halophyte roots (Barzegargolchini et al. 2017), indicating they are amongst known halophytic features to cope better under salt stress.

Our proteome and phenylpropanoid pathway analysis sought to uncover if there is a metabolic re-configuration enabling these apparent differences in root lignification capacity. We observed several processes acting together to enable this enhanced capacity. Firstly, Westonia *Nax1* and Westonia *Nax2* showed an enhancement in energy metabolism under salt stress. Significant increases of abundance occurred in glycolytic and oxidative phosphorylation enzymes (Figure 4) including fructokinase, enolase, GAPDH, PEPC, NADH-quinone oxidoreductase subunit E, ATP synthase gamma chain, cytochrome c oxidase subunit 5B, ATP synthase delta subunit and ATP synthase subunit d, which indicate the induction of respiratory metabolism under salt stress. Similarly, increases in glycolysis and respiratory metabolism was found to be involved in the enhancement of salt tolerance in cultivated barley (Wu et al. 2013). Maintenance of higher respiratory capacity is closely related to high ATP production (Hannachi et al. 2022). Increase in glycolysis and oxidative phosphorylation provides glycolytic end products and ATP for salt exclusion in order to maintain high level of salt tolerance in Westonia Nax lines. Similarly, Hannachi et al. (2022) reported that the salt tolerant Eggplant cv. Bonica used increased cellular energy production to attain sodium ion exclusion.

In addition, Westonia *Nax2* showed changes to TCA cycle function, including increases of several isoforms of the E2 acetyltranferase subunit of PDH complex, the E2 succinyltransferese subunit of 2-oxoglutarate DH complex and isocitrate dehydrogenase. However, the E1 component of PDH complex was found to decrease in abundance which suggest the differential response of mitochondrial PDC subunits under salt stress. The E1 subunit catalyses the commitment step to PDH complex catalysis (Sheng and Liu 2013). This indicates a potential sensitivity of mtPDC to salt which then could alter downstream pyruvate metabolism. Our 2-Way ANOVA genotype x treatment interaction response showed significantly lower abundance of PDC and higher abundance of malic enzyme and alanine aminotransferase in Gladius compared to Mocho and Westonia *Nax* lines indicating a potential metabolic re-arrangement downstream of pyruvate in Gladius. Inhibition of pyruvate metabolism was also reported in roots of both wild and cultivated barley in response to salt stress (Wu et al. 2013)

The proteomic data in Mocho and Westonia *Nax1* also highlighted the contribution of aromatic amino acid synthesis through shikimate pathway in response to salt stress and the lower/absence of this response in Gladius. Mocho maintained significantly higher abundance of rate limiting enzymes of shikimate pathway including DAHP synthase and 3-dehydroquinate synthases compared to Gladius, regardless of exposure to salt. Further, unique DAP response of Westonia *Nax1* showed increase of abundance of DAHP synthase, anthranilate phosphorybosyltransferase and tryptophan synthase which suggests the diversion of carbon flow through shikimate pathway to generate aromatic amino acids under salt stress. The shikimate pathway produces aromatic amino acids including tryptophan, phenylalanine and tyrosine (Maeda and Dudareva 2012). Further, shikimate pathway in plants provides an entry point to the biosynthesis of phenylpropanoids (Vogt 2010). Constitutively higher abundance of proteins involved in phenylpropanoid pathway including cinnamate 4-hydrolase (C4H), p-coumarate 3-hydrolase (C3H), and cinnamoyl alcohol dehydrogenase (CAD) provides evidence on metabolic re-arrangement in Mocho, Westonia *Nax1* and Westonia *Nax2* to have a higher capacity to use enhanced glycolytic rates for root lignification. In agreement with the proteomic changes highlighted by the 2-way ANOVA genotypic effect, the targeted metabolite analysis also showed that most of the metabolites involved in phenylpropanoid pathway were significantly more abundant in either Mocho or Westonia *Nax* lines compared to Gladius. However, most of these metabolites showed significant changes under salt treatment conditions, indicating the inducible metabolic response of roots under salt stress with respect to monolignol synthesis. Increased abundance of phenylpropanoid metabolites was also reported by Zhu et al. (2021) in roots of salt tolerant perennial legume *Sophora alopecuroides*. Further, Zhu et al. (2021) proposed that the flavonoids and lignin may participate in the removal of reactive oxygen species (ROS) in roots, thereby reducing the damage caused under salt stress.

Significantly higher abundance of coniferyl alcohol under salt stress in Mocho and Westonia *Nax2* and sinapyl alcohol in Westonia *Nax2* confirms the metabolic capability of Mocho and Westonia *Nax* lines to promote lignification under salt stress. This result was also consistent with the histochemical staining of the root cross sections in Mocho, Westonia *Nax1* and Westonia *Nax2* lines that showed more intensified lignification in the VC compared to Gladius under both control and salt treatment conditions. Further, Westonia *Nax* lines produce effective apoplastic barriers in endodermis which could potentially contribute to the low Na^+^ levels in shoots. Overall, our results show the intense lignification in the VC in roots, and effective apoplastic barrier formation in the salt tolerant wheat lines, and a series of different but complementary metabolic re-configurations enabling these root lignification capacities (Figure 6).

**Figure 6.**
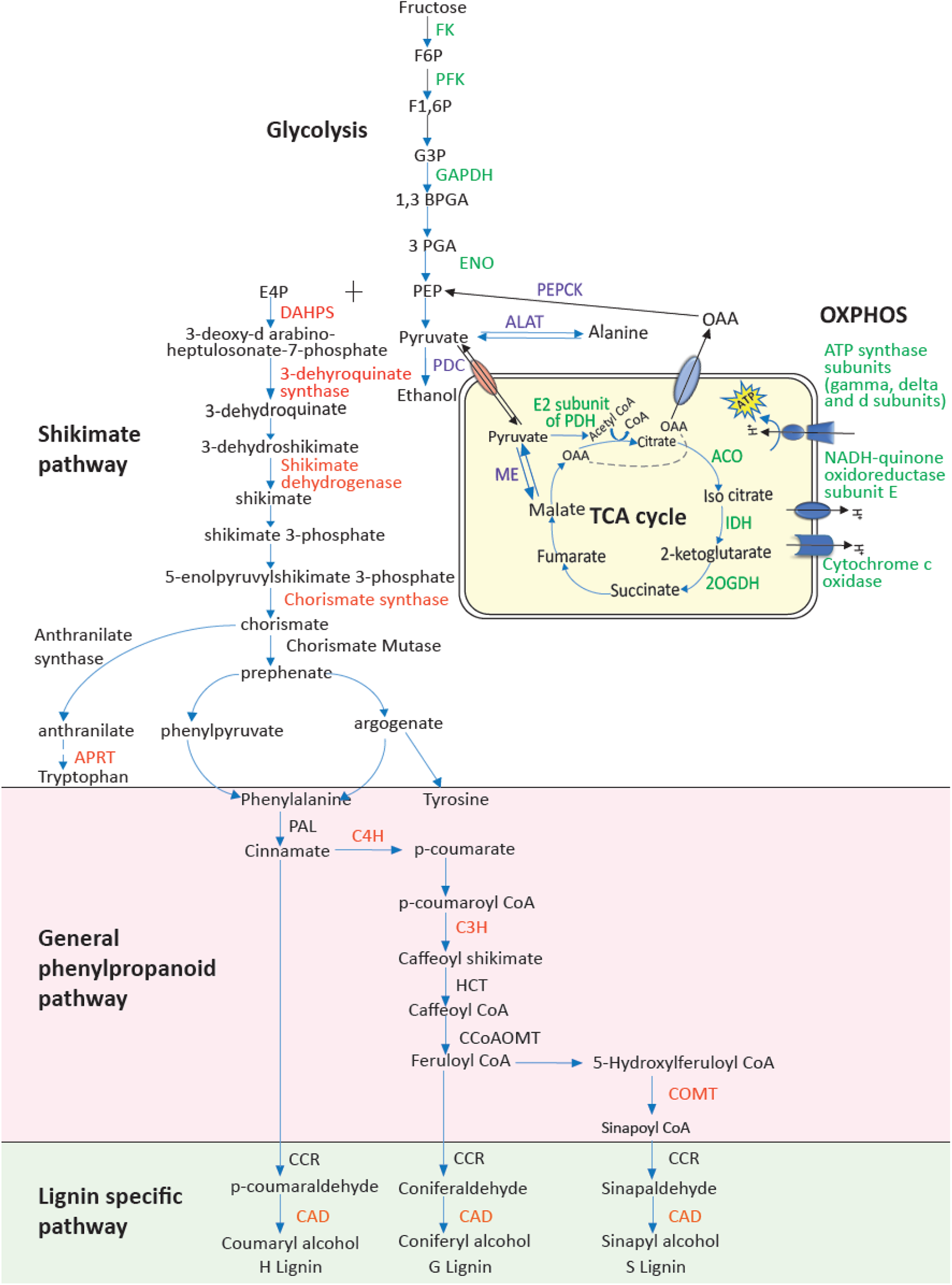
Schematic illustration of constitutive and induced proteomic changes of roots of bread wheat varieties. Green-proteins induced/increased in abundance of energy metabolism related proteins in Westonia Nax1 and Nax2 under salt stress, Red-proteins found to be more significantly abundant in Mocho and Westonia *Nax* lines compared to Gladius in 2-Way ANOVA genotype comparisons/significantly induced in Mocho or *Nax* lines under salt stress, Purple-proteins increased in abundance in Gladius under salt stress compared to Mocho and Westonia *Nax* lines in 2-WAY ANOVA genotype x treatment comparisons (FK-Fructokinase, PFK-Phosphofructokinase, GAPDH-Glyceraldehyde 3-phosphate dehydrogenase, ENO-Enolase, ACO-Aconitase, IDH-Isocitrate dehydrogenase, 2OGDH-2-oxoglutarate dehydrogenase, PDH-Pyruvate dehydrogenase, PDC-Pyruvate dehydrogenase complex, ALAT-Alanine aminotransferase, PEPCK-Phosphoenolpyruvate carboxykinase, ME-Malic Enzyme, OAA- Oxaloacetate, DAHPS-3-deoxy-d-arabino-2-heptulosonate 7-phosphate synthase, ESPS synthase-5-enolpyruvylshikimate-3-phosphate synthase, PDT-Prephenate dehydratase, ADT-Arogenate dehydratase, APRT-Anthranilate phosphoribosyltransferase, PAL-Phenylalanine ammonia lyase, C4H-Cinnamic acid4 hydroxylase, C3H-p-coumaroyl shikimate3ʹ hydroxylase, HCT-Hydroxycinnamoyl-coenzyme A shikimate:quinate hydroxycinnamoyl-transferase, CCoAOMT-Caffeoyl-C oAO methyltransferase, COMT – Caffeic acid /5-hydroxyferulic acid O-methyltransferase, CCR-Cinnamoyl CoA reductase, CAD – Cinnamoyl alcohol dehydrogenase).

## Conclusion

Here we compile evidence that the metabolic capacity in roots of bread wheat varieties contribute to salt tolerant mechanisms to cope better under salt stress through enhancing energy metabolism and primary metabolism for increased lignification. We conclude that maintenance of constitutively higher abundance of proteins involved in lignin biosynthesis and induced abundance of phenylpropanoid pathway intermediates allow Mocho, Westonia *Nax1* and Westonia *Nax2* lines to enhance early and late metaxylem lignification and formation of effective apoplastic root barriers leading to tolerance in early stages of salt stress. Better understanding of distinct salt tolerant mechanisms in roots of different wheat varieties will help develop more salt tolerant wheat to maintain yield despite the predicted salinization of agricultural land.

## Acknowledgement

BMD was the recipient of a Scholarship funded by the Grains Research and Development Corporation (UWA00173). This work was supported through funding by the Australian Research Council (ARC) (CE140100008) to AHM and ARC Future Fellowship (FT170100195) to KR. Peptide quantitation in this work was performed as a service by the WA Proteomics Facility as a node of Proteomics Australia, supported by infrastructure funding from the Western Australian State Government in partnership with Bioplatforms Australia under the Commonwealth Government National Collaborative Research Infrastructure Strategy.

## Author Contributions

BMD planned and designed research, performed the experiments, undertook data analysis and interpretation, and wrote the manuscript. KR designed experiments and participated in histochemical staining of roots and quantification of total lignin. TWR developed and conducted the targeted LCMS analysis. CS, AHM, RM and NLT participated in planning and designing the research, provided advice on the data analysis and interpretation. All the authors contributed to writing the manuscript.

## List of Supplementary Figures

Supplementary Figure 1. Differences in root growth, Na^+^ and K^+^ ion concentrations in roots after 3 days of exposure to 150 mM NaCl. (A) Root Dry weight, (B1) Total root length, (B2) Changes of root lengths that belongs to different diameter classes, (C) Na^+^ ion concentration, (D) K+ ion concentration, (E) The K^+^/Na^+^ ratio (Different letters denote significant differences obtained through Tukey’s HSD test, error bars indicate the standard error of mean, n=4).

Supplementary Figure 2. Lignin staining in the vascular cylinder (VC) of roots gown under control condition; At 75 % from the total length of Zone-I, orange/red staining in the early metaxylem of roots grown under control condition: (A) Faint orange staining in the VC of Gladius, (B) Bright orange/red staining in the VC of Mocho, (C) Bright orange/red staining in the VC of Westonia *Nax1*, and (CD) Bright orange/red staining in the VC of Westonia *Nax2*. (A-D) The “U” shaped tertiary cell walls in the endodermis with faint orange stains (arrows) in roots.

Supplementary Figure 3. Protein abundance and corresponding pathways of common DAPs in roots of four wheat varieties after 3 days of salt stress. (A) Heat map showing the abundance change (Log2 Fold change) of DAPs, (B) Pie chart showing the metabolic pathways of proteins which increased in abundance under salt stress (% values indicate the percentage of proteins which increased in abundance in each functional category compared to the total number of proteins that increased abundance, (C) Pie chart showing the pathways of proteins which decreased in abundance under salt stress (% values indicate the percentage of proteins which decreased in abundance in each functional category compared to the total number of proteins that decreased abundance)

Abbreviations: W.*Nax1*; Westonia *Nax1*, W.*Nax2*; Westonia *Nax2*.

Supplementary Figure 4. Protein abundance and corresponding pathways of common DAPs in roots of four wheat varieties after 6 days of salt stress. (A) Heat map showing the abundance change (Log2 Fold change) of DAPs, (B) Pie chart showing the metabolic pathways of proteins which increased in abundance under salt stress (% values indicate the percentage of proteins which increased in abundance in each functional category compared to the total number of proteins that increased abundance, (C) Pie chart showing the pathways of proteins which decreased in abundance under salt stress (% values indicate the percentage of proteins which decreased in abundance in each functional category compared to the total number of proteins that decreased abundance)

Abbreviations: W.*Nax1*; Westonia *Nax1*, W.*Nax2*; Westonia *Nax2*.

## Notes

### Competing Interest Statement

The authors have declared no competing interest.

