## Supplementary Figures 1,2,3,4 for "Metabolic adaptations leading to lignification in wheat roots under salinity stress"

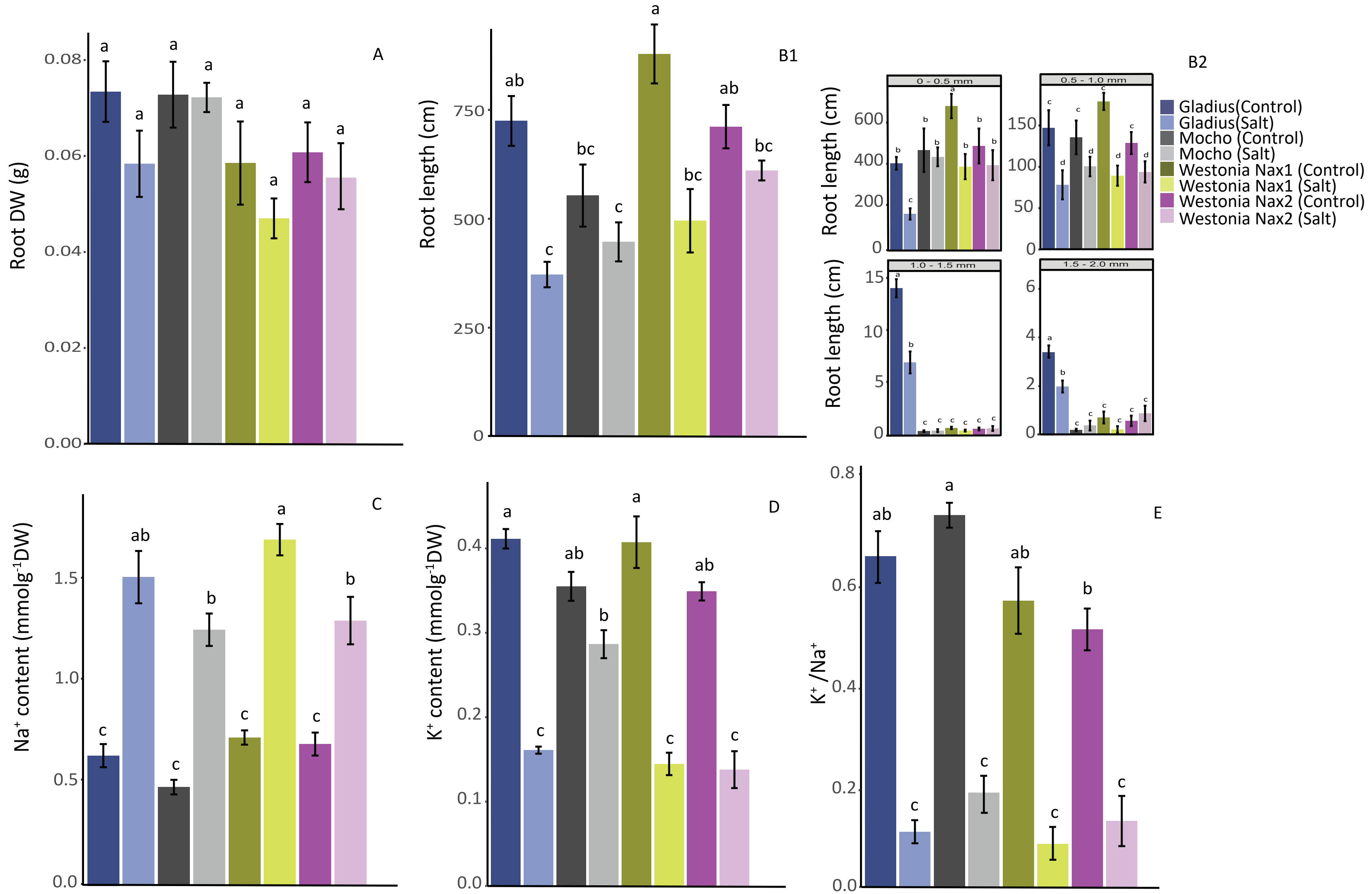


Supplementary Figure 1. Differences in root growth, $Na^{+}$ and $K^{+}$ ion concentrations in roots after 3 days of exposure to 150 mM NaCl. (A) Root Dry weight, (B1) Total root length, (B2) Changes of root lengths that belongs to different diameter classes, (C) $Na^{+}$ ion concentration, (D) K+ ion concentration, (E) The K+/Na+ ratio (Different letters denote significant differences obtained through Tukey's HSD test, error bars indicate the standard error of mean, n=4)

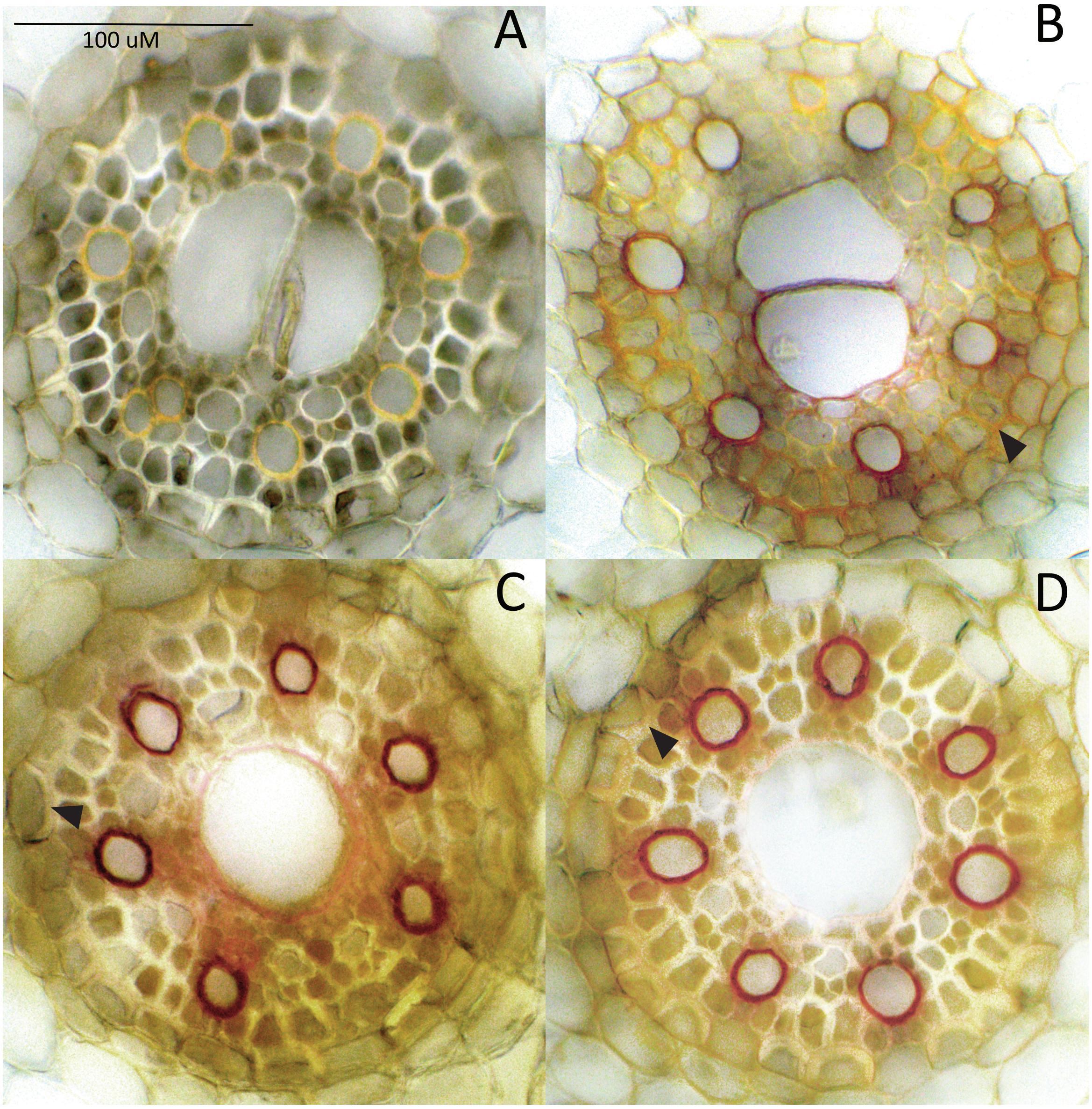


Supplementary Figure 2. Lignin staining in the vascular cylinder (VC) of roots gown under control condition; At 75 % from the total length of Zone-I, orange/red staining in the early metaxylem of roots grown under control condition: (A) Faint orange staining in the VC of Gladius, (B) Bright orange/red staining in the VC of Mocho, (C) Bright orange/red staining in the VC of Westonia *Nax1*, and (CD) Bright orange/red staining in the VC of Westonia *Nax2*. (A-D) The “U” shaped tertiary cell walls in the endodermis with faint orange stains (arrows) in roots.

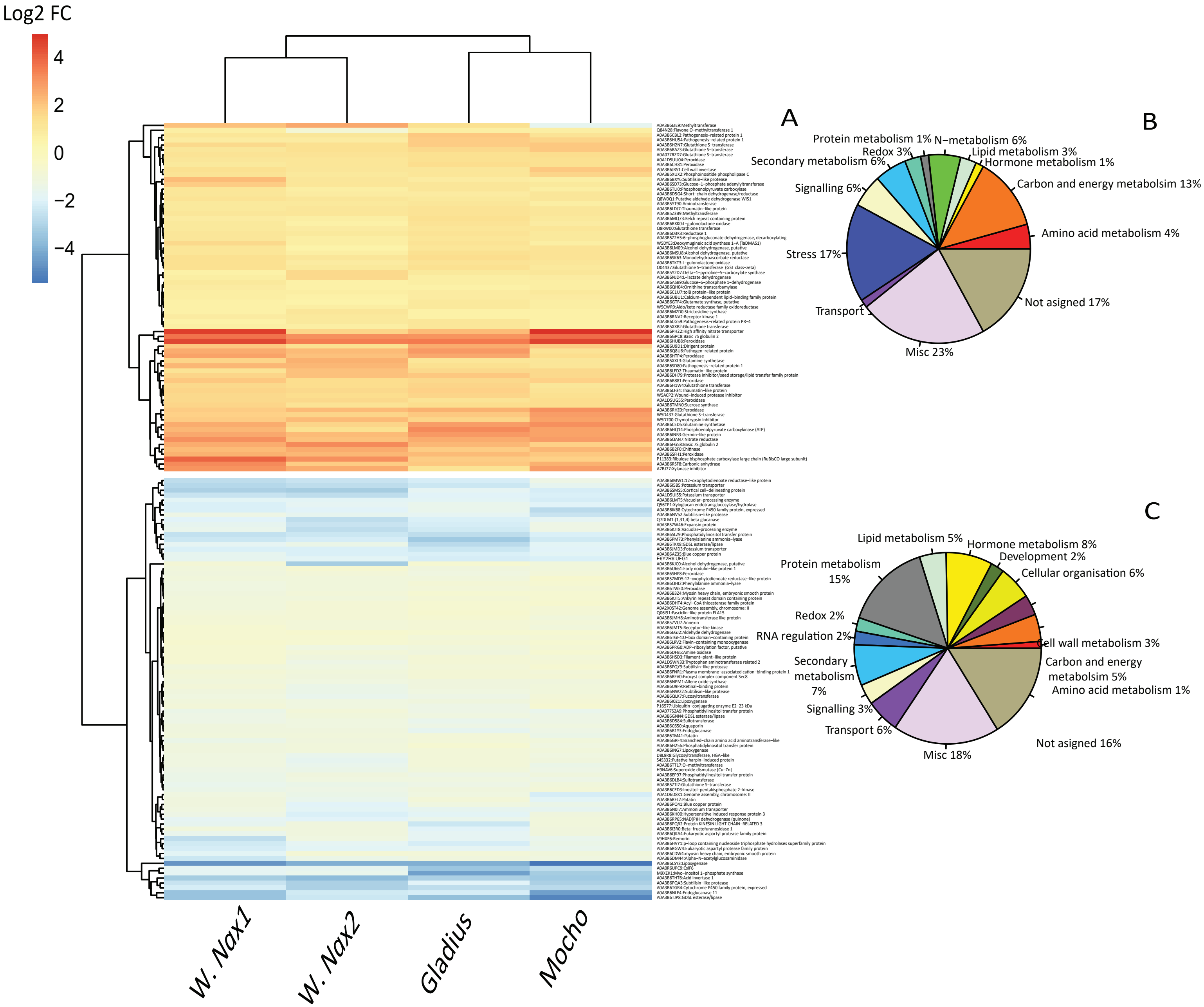


Supplementary Figure 3. Protein abundance and corresponding pathways of common DAPs in roots of four wheat varieties after 3 days of salt stress. (A) Heat map showing the abundance change (Log2 Fold change) of DAPs, (B) Pie chart showing the metabolic pathways of proteins which increased in abundance under salt stress (% values indicate the percentage of proteins which increased in abundance in each functional category compared to the total number of proteins that increased abundance, (C) Pie chart showing the pathways of proteins which decreased in abundance under salt stress (% values indicate the percentage of proteins which decreased in abundance in each functional category compared to the total number of proteins that decreased abundance)

Abbreviations: W.*Nax1;* Westonia *Nax1*, W.*Nax2*; Westonia *Nax2*.

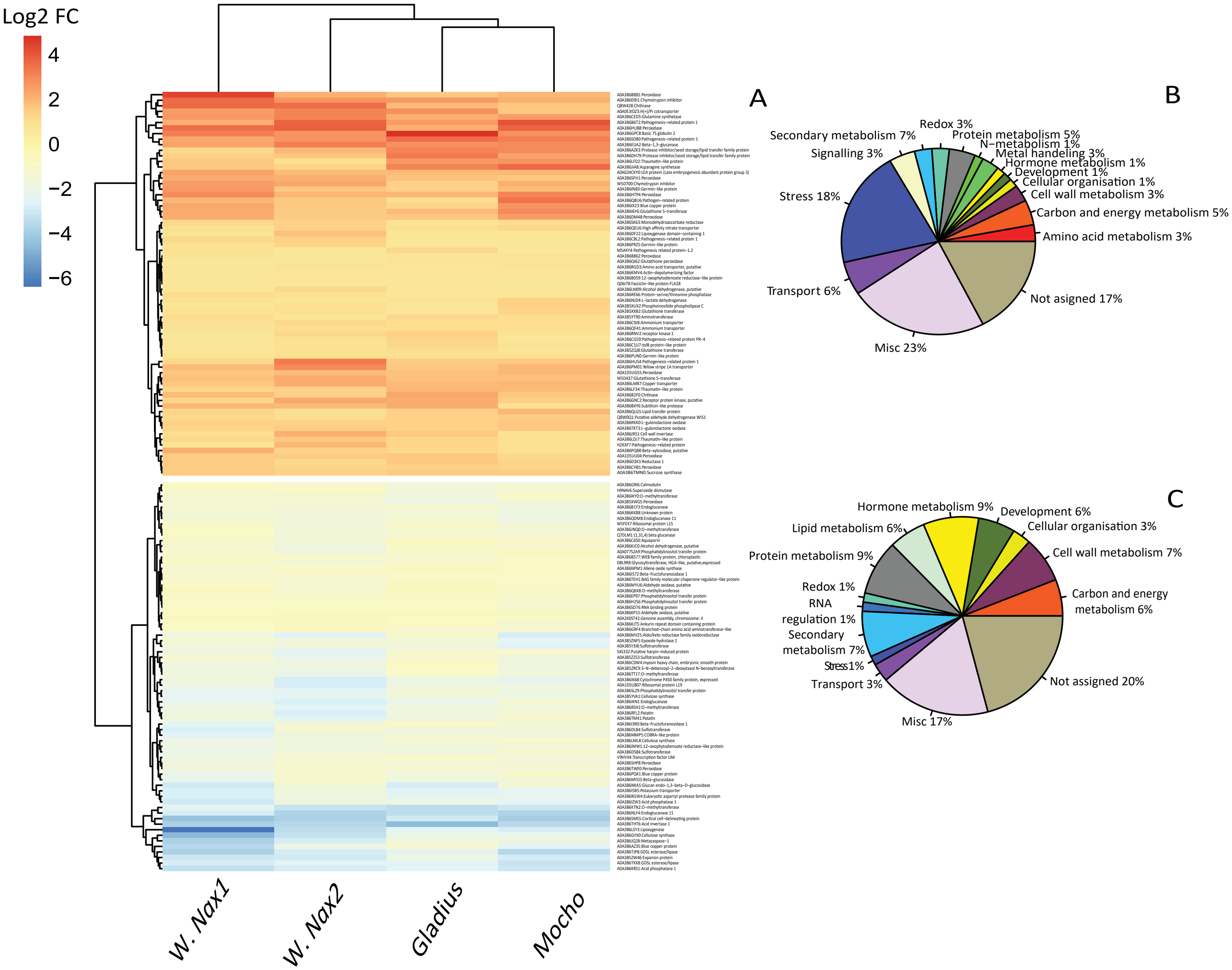


Supplementary Figure 4. Protein abundance and corresponding pathways of common DAPs in roots of four wheat varieties after 6 days of salt stress. (A) Heat map showing the abundance change (Log2 Fold change) of DAPs, (B) Pie chart showing the metabolic pathways of proteins which increased in abundance under salt stress (% values indicate the percentage of proteins which increased in abundance in each functional category compared to the total number of proteins that increased abundance, (C) Pie chart showing the pathways of proteins which decreased in abundance under salt stress (% values indicate the percentage of proteins which decreased in abundance in each functional category compared to the total number of proteins that decreased abundance)

Abbreviations: W.*Nax1*; Westonia *Nax1*, W.*Nax2*; Westonia *Nax2*.
